# Expanded Analysis of Pigmentation Genetics in UK Biobank

**DOI:** 10.1101/2022.01.30.478418

**Authors:** Erola Pairo-Castineira, Jaime Cornelissen, Konrad Rawlik, Oriol Canela-Xandri, Stacie K. Loftus, William J. Pavan, Kevin M. Brown, Albert Tenesa, Ian J. Jackson

**Affiliations:** MRC Human Genetics Unit, Institute of Genetics and Cancer, University of Edinburgh, EH4 2XU, UK; Roslin Institute, University of Edinburgh, Easter Bush, EH25 9RG, UK; National Human Genome Research Institute, National Institutes of Health, Bethesda, MD 20892-2152, USA; Division of Cancer Epidemiology and Genetics, National Cancer Institute, National Institutes of Health, Bethesda, MD 20892, USA

## Abstract

The genetics of pigmentation is an excellent model for understanding gene interactions in a trait almost entirely unaffected by environment. We have analysed pigmentation phenotypes in UK Biobank using DISSECT, a tool which enables genome-wide association studies (GWAS) whilst accounting for relatedness between individuals, and thus allows a much larger cohort to be studied. We have increased the number of candidate genes associated with red and blonde hair colour, basal skin colour and tanning response to UV radiation. As previously described, we find almost all red hair individuals have two variant *MC1R* alleles; exome sequence data expands the number of associated coding variants. Rare red-headed individuals with only a single *MC1R* variant are enriched for an associated eQTL at the *ASIP* gene. We find that females are most likely to self-report red or blonde hair, paler skin and less tanning ability than men, and that variants at *KITLG, MC1R, OCA2* and *IRF4* show significant sex differences in effect. After taking sex into account, pigmentation phenotypes are not correlated with sex hormone levels, except for tanning ability, which shows a positive correlation with testosterone in men. Across the UK there is a correlation between place of birth and hair colour; red hair being more common in the north and west, whilst blonde hair is more common in the east. Combining GWAS with transcriptome data to generate a transcriptome wide association study identifies candidate genes whose expression in skin or melanocytes shows association with pigmentation phenotypes. A comparison of candidates associated with different pigmentation phenotypes finds that candidates for blonde hair, but not skin colour, are enriched for skin and hair genes suggesting that it may be hair shape and structure that impacts hair colour, rather than the melanocyte/keratinocyte interaction.

## Introduction

The basal pigmentation of human skin and hair is a polygenic trait on which environment has little impact. The tanning response of skin to the environmental insult of UV radiation (sunlight) also has a strong genetic basis. All three traits are correlated, although the correlation is far from absolute. Typically, individuals with dark hair have darker skin, have less propensity to burn in strong sunlight and tan well, but there are others with dark hair who have pale skin which easily burns.

The strong genetic underpinnings of pigmentation make it an excellent model for understanding complex genetic phenotypes. Identifying those genes that affect more than one pigmentary phenotype compared to those unique to a single trait casts light on the biology of the melanocytes, which produce the pigment and of the keratinocytes of the skin and hair with which they interact and transfer pigment. Furthermore, the strong causal link between pale skin and a poor tanning response and skin cancer highlights the medical importance of understanding the genetics of skin pigmentation and response to UV.

The genetics of human pigmentation has been reviewed by Pavan and Sturm ^1^. Genetic variation in hair colour has been analysed in relatively homogenous populations, almost exclusively of European-origin ^2–8^. On the other hand, the genetics of variation in skin pigmentation has been studied not only in Europeans^9,10^ but also in South Asians^11^, Africans^12^ and in admixed African, European and Native American populations^13–16^. The tanning response is partially correlated with skin pigmentation, and association studies have identified loci in common, but there are also tanning loci not associated with skin colour^17,18^.

UK Biobank (UKB) is a very large population wide survey of phenotypic and clinical variation linked to genotype data^19^. Previous work by ourselves and others have mapped genetic associations in UKB with hair pigmentation and tanning response^7,8,18^. These studies have used a subset of UKB subjects, in part because relatedness between individuals confounds the analysis. Using DISSECT, an algorithm we have developed that can take into account relatedness ^20^, we have expanded the GWAS of hair colour in UKB, and have extended the associations to include skin colour and tanning response to UVR.

In general GWAS identify numerous associated SNPs at any locus due to linkage disequilibrium. An alternative method, transcriptome wide association study, TWAS, uses transcriptome-wide expression data coupled with genotypes to predict gene expression associations in a GWAS cohort. Furthermore, most SNP associations are not within coding regions and it is likely that non-coding, causative, SNPS affect gene expression. TWAS may thus directly identify the gene affected, if not the SNP that causes the altered gene expression^21^.

## Results

Our previously published analysis of hair colour^8^ used a logistic model on ∼340,000 unrelated white British individuals in UK Biobank. We excluded all but one member of any group of related individuals (identified by genotype), which would otherwise introduce a confounder for the genetic relationship between individuals in the analysis. We also excluded from this analysis white non-British participants. Overall this resulted in ∼150,000 participants being excluded. We used a tool we have previously developed called DISSECT^20^ which performs mixed linear models in a fast and scalable way, and thus permits inclusion of those previously excluded. This allows analysis of a UKB cohort of much increased size, giving correspondingly increased power to detect associations. Table 1 shows the pigmentary phenotypes of the entire analysed cohort.

**Table 1:** Self-reported phenotypic data for UK Biobank European individuals

| TRAIT | VALUE | EUROPEANS | % | MALE | % | FEMALE | % |
| --- | --- | --- | --- | --- | --- | --- | --- |
| Hair Colour | Blonde | 53100 | 11.74% | 20247 | 9.79% | 30809 | 12.55% |
|  | Red | 21451 | 4.74% | 7845 | 3.79% | 12708 | 5.18% |
|  | Light brown | 191665 | 42.39% | 80939 | 39.15% | 102841 | 41.90% |
|  | Dark brown | 177479 | 39.25% | 77134 | 37.31% | 92851 | 37.83% |
|  | Black | 21244 | 4.69% | 16276 | 7.87% | 4114 | 1.67% |
|  | NA | 6398 | 1.41% | 4289 | 2.07% | 2109 | 0.86% |
| Skin Colour | Very Fair | 36457 | 8.06% | 13575 | 6.57% | 22882 | 9.32% |
|  | Fair | 319466 | 70.65% | 148966 | 72.05% | 170500 | 69.46% |
|  | Light Olive | 81824 | 18.10% | 35310 | 17.08% | 46514 | 18.95% |
|  | Dark Olive | 6969 | 1.54% | 4318 | 2.09% | 2651 | 1.08% |
|  | Brown | 1864 | 0.41% | 1424 | 0.68% | 440 | 0.18% |
|  | Black | 31 | 0.006% | 20 | 0.009% | 11 | 0.004 |
|  | NA | 5550 | 1.22% | 3116 | 1.50% | 2434 | 0.99% |
| Ease of Skin tanning | Very tanned | 91094 | 20.14% | 53077 | 25.67% | 38017 | 15.48% |
|  | Moderately tanned | 178559 | 39.49% | 85509 | 41.36% | 93050 | 37.91% |
|  | Midly or occasionally tanned | 95005 | 21.01% | 34910 | 16.89% | 60095 | 24.48% |
|  | Never tan | 78392 | 17.33% | 29720 | 14.38% | 48672 | 19.83% |
|  | Only burn | 0 | 0 | 0 | 0 | 0 | 0 |
|  | NA | 9111 | 2.015% | 3513 | 1.69% | 5598 | 2.28% |

### Red Hair

We performed a GWAS of ∼450,000 European individuals in UK Biobank, comparing individuals with red hair versus black or brown hair. This identified 655 independent associated SNPs (Figure 1a) (accounting for 35% heritability). Comparing these GWAS results with those obtained previously using only unrelated white British we find that there is an increase in significant SNPs of about 24% (10,797 compared to 8,716) (Supplementary Figure 1a),. By contrast the number of SNPs that lose significance when using data from all Europeans data is only 18 (0.2%).

**Figure 1:**
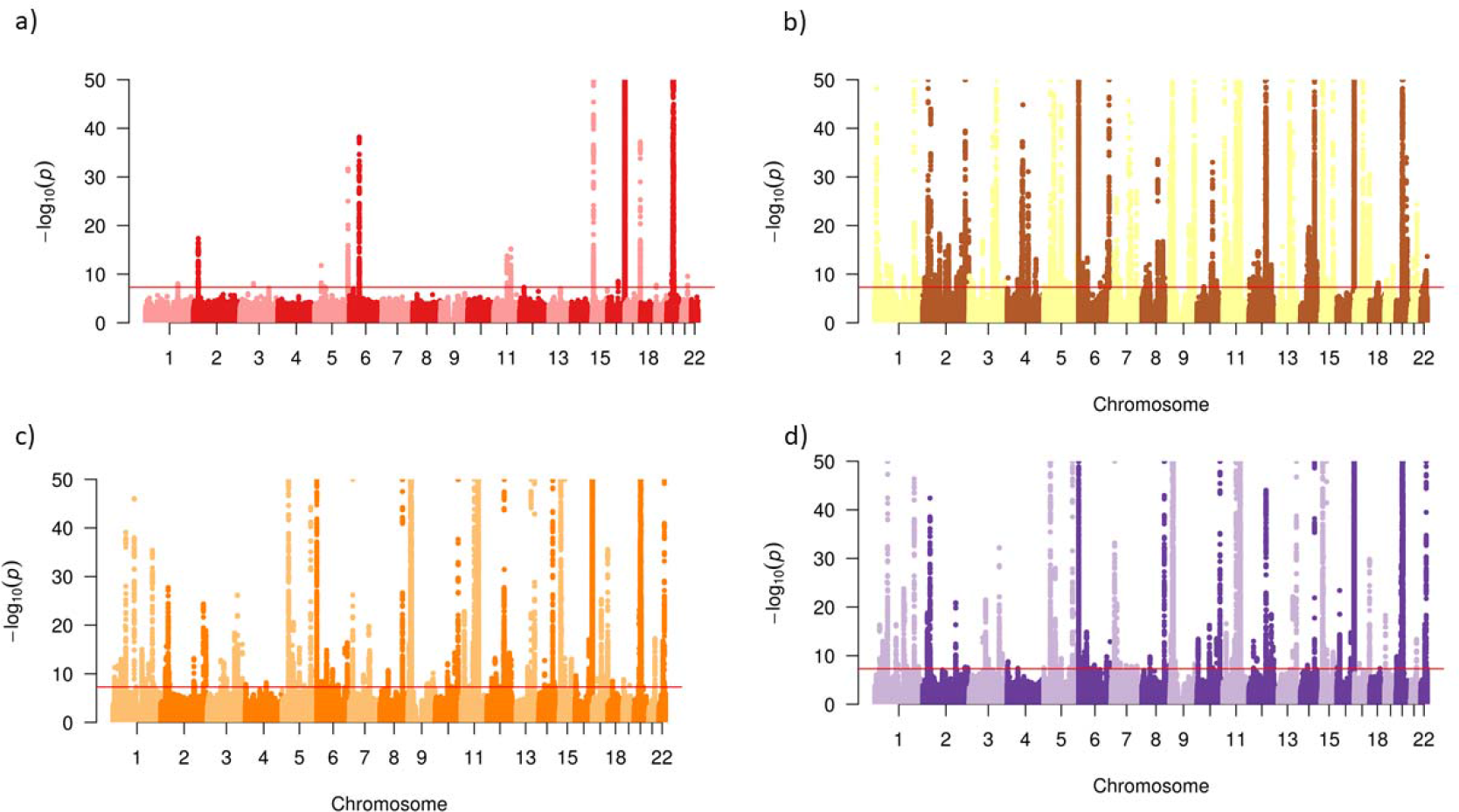
Manhattan Plots. Manhattan plots for GWAS using European UK Biobank individuals for a) Red vs Brown and Black hair, b) Blonde vs Brown and black hair c) Skin Colour and d) Ease of skin tanning.

To identify candidate genes affected by the significant associated SNPs we looked at the closest gene or genes to the 772 lead SNPs, identifying 128 putative causal genes (Supplementary Table 1). Of these, 48 are on chromosome 16, in the region of *MC1R*, highlighting the complex LD pattern and gene expression correlation in the region (Supplementary Figure 2 and see below). A further 50 genes are on chromosome 20, surrounding *ASIP*, a gene also strongly associated with red hair. These 50 genes may be identified by virtue of weak LD with the strong signal at *ASIP*. If these loci are excluded as candidates then only an additional 31 remain. Furthermore, at 18 of these loci the associated SNP lies between two genes; both are identified as candidates but likely only one will be the causative gene. In addition, we identify both *OCA2* and *HERC2* in this analysis, but work by others has shown that variants in *HERC2* are affecting transcription of *OCA2*, which is the likely causative gene. Thus we identify 21 gene loci as associated with red hair (Supplementary Table 2)

*MC1R* is well known as the gene at which almost all red-hair individuals have two variant alleles. Exome sequence data is available for ∼200K UK Biobank individuals, and we interrogated this data for *MC1R* variants in the 8,496 individuals with red hair (Supplementary Table 3 shows hair colour phenotype data from the 200K individuals with WES data available). We found 106 *MC1R* protein coding variants (missense, frameshift or gain of stop codon) in red hair individuals, of whom 96.2% had 2 variants (Supplementary Table 4). From the 106 protein coding variants, 99 of them have an assigned rsid, while 7 are variants not previously described. From these 106 variants, 45 are only present in individuals that also have 2 additional, previously described *MC1R* variants. We suggest that these 45 variants do not affect hair colour, leaving 61 red-hair associated *MC1R* coding variants and only 13 individuals who have 3 functional *MC1R* variants. Supplementary Table 5 shows the 61 *MC1R* coding variants associated with red hair across the entire WES cohort.

The reported red hair for the subjects with only 0 or 1 *MC1R* variants may be due to an error in self-reported hair colour, non-coding red hair variants not detected by exome sequencing, a dominant action of certain *MC1R* variants in combination with other genes, or indeed rare non-*MC1R* associated red hair. To check if variants at genes other than *MC1R* could suffice to produce red hair, in combination with a single *MC1R* variant, we took the 2.4% red-hair individuals (207 individuals) with 1 *MC1R* coding variant and looked at enrichment for the *ASIP* risk allele in the *ASIP* eQTL (rs6059655)^8^ the next-most strongly associated locus. We found that 55% of them have the *ASIP* eQTL risk allele, compared to 31% of the total individuals with red hair, which is a significant enrichment over the ones that have 2 *MC1R* coding variants (Fisher exact test p-value=6.6×10^−11^). There is enrichment in this cohort for heterozygotes for the ASIP variants (Fisher exact test P=7.02×10^−6^) as well as homozygotes (Fisher exact test P=1.82×10^−6^). Our previous work has demonstrated epistasis between weak red hair alleles of *MC1R* and variation at *ASIP*. Here we suggest that in certain genetic backgrounds heterozygosity for *MC1R* variants may be sufficient for red hair, as previously also suggested by Duffy et al^22^.

We have previously described an association between a variant 120bp 5’ of the transcriptional start site of *MC1R* (rs3212379) strongly associated with red hair^8^. Examination of WES data reveals that there is a coding variant, rs796296176, which is a 1bp insertion at base 85 of the cds and consequently introduces a frameshift at codon 29 which is in high ld (r^2^ ∼0.8) with rs3212379. From the individuals with the rs3212379 variant, WES data available and red hair, 376/380 have the frameshift variant, and the other 4 have 2 *MC1R* coding variants. This frameshift variant has not been associated with red hair before and it was not genotyped or imputed in our previous analysis, but it is most likely the functional variant rather than the 5’ variant.

Sequence analysis of rare red-haired individuals in Jamaica found several alleles seen in European populations, presumably present by introgression^23^, but also found the stop-gained mutation, rs201533137 at codon 23, in combination with these alleles in some individuals. This variant has only been seen in African populations to date, at frequencies up to 1.5%. We find 45 of the 200K exome sequenced individuals are carriers of this allele. None of them have this allele in combination with other *MC1R* variants and none have red hair. However, all of them by self-report and by genotype are of African, Caribbean or mixed European/Caribbean/African descent. We suggest that homozygotes for this variant will not be uncommon in certain African populations.

#### Blonde Hair

We analysed with DISSECT blonde vs brown and black hair for the ∼450,000 subjects of European descent in UK Biobank. We found 1337 independent SNPs related to blonde hair (Supplementary Table 1). Comparing the blonde hair colour GWAS to our previous study, with unrelated white British, we found a 45% increase in associated SNPs from 22,419 to 32,349, and just 201 SNPs (0.9%) lose significance in this new analysis. As in the red hair colour GWAS, we can see the additional ∼100,000 individuals enable the identification of additional associations (Supplementary Figure 1b). Applying the same criteria as we have done for red hair to the 1,777 lead SNPs, excluding 34 associations in LD at *MC1R* and 29 in LD with *ASIP* we identify 408 candidate genes for blonde hair (Supplementary Table 2). Again, many of these are pairs associated with a single intragenic SNP. Accounting for these, and the *HERC2*/*OCA2* complex, leaves a maximum of 344 associated gene loci, an increase from 205 in our previous analysis. However, there are other genomic regions, around strongly blonde-hair associated SNPs that may also show association through weak LD and hence reduce this number.

### Skin Pigmentation and Tanning Response

Skin pigmentation in UK Biobank is self-reported data and coded with a scale where 1=very fair, 2=fair, 3=light olive, 4=dark olive, 5=brown, 6=black and -1, -3 = NA. Treating the phenotype as a numerical trait we performed a skin pigmentation GWAS using DISSECT for ∼450,000 European origin individuals. We found 1546 independent SNPs (Figure 1c). Curating the list in the same way as in the hair analysis results in 337 genes. Removing 44 genes around *MC1R* and 64 around *ASIP* leaves 229, of which 40 were pairs identified by a single intergenic SNP. Counting only one of these as well as excluding *HERC2* leaves a maximum of 188 skin colour associated genes (Supplementary Table 2).

The Visigen consortium performed a skin colour meta-analysis of GWAS from different European cohorts (n=17,262) finding 388 significant SNPs^9^. All significant genes in their study are also significant in our analysis, and we increase the number of genes up to more than 30 fold from 6 (*SLC24A5, IRF4, OCA2, HERC2, MC1R, ASIP*) to 188

Self-reported data of ease of skin tanning in UK Biobank is coded as 1=get very tanned, 2=get moderately tanned, 3=get mildly or occasionally tanned, 4=never tan, 5=only burn. -1,-3=NA. Again treating the phenotype as a numerical trait we performed a GWAS using the European origin population (∼450,000 individuals), finding 2,010 lead SNPs in 292 genes (Figure 1d). Editing the lists as before leaves 179 genes, identified by 155 SNPs resulting in a maximum of 154 associated genes (Supplementary Table 2).

We compared these results with those published by Visconti et al.^18^, who performed a GWAS for ease of tanning using two groups (easily tan=groups 1,2, and 3 vs burn=groups 4 and 5) using the first phase of UK Biobank. Comparing the 10,834 significant SNPs (p-val<5×10^−8^) found by them, we find 8,902 pass quality control in our analysis, and only 9 are not significant. In addition our analysis finds 19,835 additional significant SNPs.

### Sex Differences in Pigmentation

We find phenotypic differences between sexes in pigmentation traits between UK Biobank European individuals which consists of 206,729 men and 245,432 women, shown in Table 1. Women tend to have lighter skin and hair than men, although as UK Biobank is self-reported data, it is difficult to differentiate between self-reporting bias and true differences in pigmentation phenotypes.

First, we calculated the Pearson correlation between the self-reported data and sex to see if differences are significant. The phenotypic correlation between skin colour and sex is 0.021 (p-value=10^−44)^, for ease of skin tanning the correlation is 0.122 (p-value <10^−308^), for red hair the correlation is 0.040 (p-value=10^−157^), and for blonde hair the correlation is 0.046 (p-value=10^−214^).

We then performed a sex-specific GWAS for each pigmentation phenotype (skin colour, ease of tanning, red hair and blonde hair) using DISSECT and calculated the genetic correlation of between sexes for each phenotype with LDscore regression^24^, showing that correlation between sex-specific GWAS is high, but not one (except for blonde hair). The lowest genetic correlation is in red hair with a correlation of 0.95 (Supplementary Table 6). Comparisons of SNP effects between sex-specific GWAS is shown in Figure 2a-d where the effect of each SNP is shown by men and women. We performed a t-test in order to find SNPs with significant effect differences between men and women in pigmentation traits (Supplementary Tables 7-10). For red hair we found significant differences only on chromosome 16, the *MC1R* region, and on chromosome 20 near ASIP. In blonde hair, SNPs with significant differences are near *KITLG, MC1R, OCA2* and *IRF4*. In ease of skin tanning at *IRF4, OCA2* and *MC1R* and in skin colour, apart from a single SNP on chromosome 1, differences are found only at *IRF4*.

**Figure 2:**
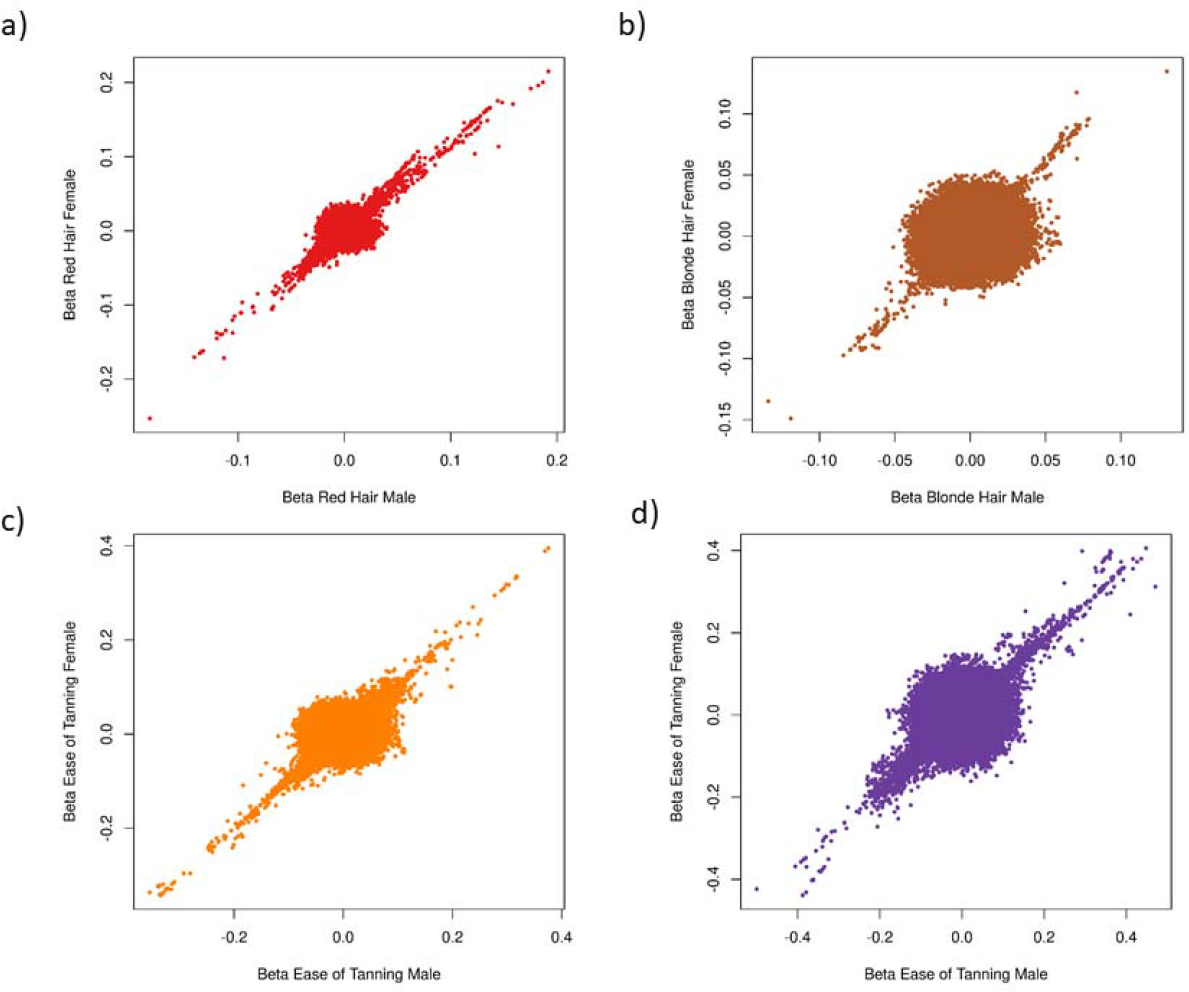
Comparison of effects between males and females. Beta comparison between a male-specific GWAS and a female-specific GWAS for a) Red Hair b) Blonde Hair c) Skin Colour and d) Ease of skin tanning

We asked if sex hormones affected pigmentation traits using estradiol and testosterone levels available in UK Biobank stratified by sex (Supplementary Figure 3). Estradiol levels are available for 68,880 individuals and testosterone levels for 390,685. Correlation between pigmentation phenotypes and sex-stratified hormone levels (Supplementary Table 11) are not significant, except for ease of tanning and testosterone levels in males (p-value<0.05). When Polygenic Risk Scores (PRS) for each pigmentation trait are used instead of the self-reported phenotype, we did not find any significant correlation after multiple test correction (Supplementary Table 12).

Estradiol levels in women change after menopause, which could reduce the correlation between the traits and hormone levels, since UK Biobank hormone levels are measured in many instances after menopause. We repeated analysis of the correlation between pigmentation phenotypes and estradiol levels using 26,426 women who had estradiol level measurements available and had not undergone menopause at the age of recruitment, without finding significant correlation between estradiol levels and pigmentation traits (p-value red hair=0.43, p-value blonde hair=0.7, p-value skin colour =0.76 and p-value ease of tanning=0.65).

### Geographical Distribution of Traits and GRS

UKB records the geographical coordinates of the birthplace of each participant. We examined this data and find a geographical correlation between pigmentation traits and North-South and East-West birthplace coordinates in the UK. We find individuals in the North tend to have more red hair, fairer skin, and tan less and burn more than individuals in the south. The same is true for people born in the west, while people born in the east have more blonde hair (Supplementary Table 13).

To confirm these trends and to mitigate self-reporting bias, we used PRS to calculate the correlation between each trait and N-S and E-W coordinates, finding again a significant correlation with all traits and both N-S and E-W (Supplementary Table 13). However, for red hair and N-S coordinates, the correlation using PRS is significant but lower than the correlation using the self-reported phenotype, which may suggest that in self-reported data people born in the north are more likely to report that they have red hair than in the south.

### Transcriptome-Wide Association Studies

We performed a transcriptome-wide association study (TWAS) for pigmentary phenotypes in 3 different datasets: sun-exposed skin and not sun-exposed skin from GTex^25^ and melanocyte cell lines^26^.

The analysis identifies more genes using the GTex data, reflecting the larger number of samples, and hence greater power, in this dataset. In all we find, combined across all datasets, 82 genes associated with red hair, 276 with blonde hair, 220 with skin colour and 188 with ease of tanning (Supplementary Tables 14-17).

As most GWAS significant SNPs are not within the coding region of genes we anticipate that the phenotypic effect of these variants is via modulation of gene expression. To further refine our list of candidate genes we determined which of our curated list of GWAS candidates were also found in the TWAS (Supplementary Table 18). This will exclude any associated coding variants that do not have an effect on mRNA levels. Following this filtering step, we find 7 genes associated with red hair, 72 with blonde hair, 53 with skin colour and 42 with tanning ability. Only 3 genes are in all 4 of these categories; *OCA2, ASIP* and, surprisingly, *MC1R*. All characterised associations of *MC1R* with pigmentation are non-synonymous, frameshift or stop-gained variants, which are not expected have an effect on mRNA expression. Many genes in the region of *MC1R* on chromosome 16 are found in the TWASs. We performed a Bayesian analysis, to highlight the probabilities of causal genes in this region of the genome. The Posterior probability for causality is similar in all genes showing the complexity of associations in this region, as previously highlighted^8^ (Supplementary Figure 4).

### Genes in Common and Unique to Traits

We calculated the genetic correlation between the different pigmentation traits using LD score^27^. The highest genetic correlation is between skin colour and ease of skin tanning (cor=0.88), while the genetic correlation between blonde hair and red hair is just 0.36. The correlation between the traits is shown in Figure 3.

**Figure 3:**
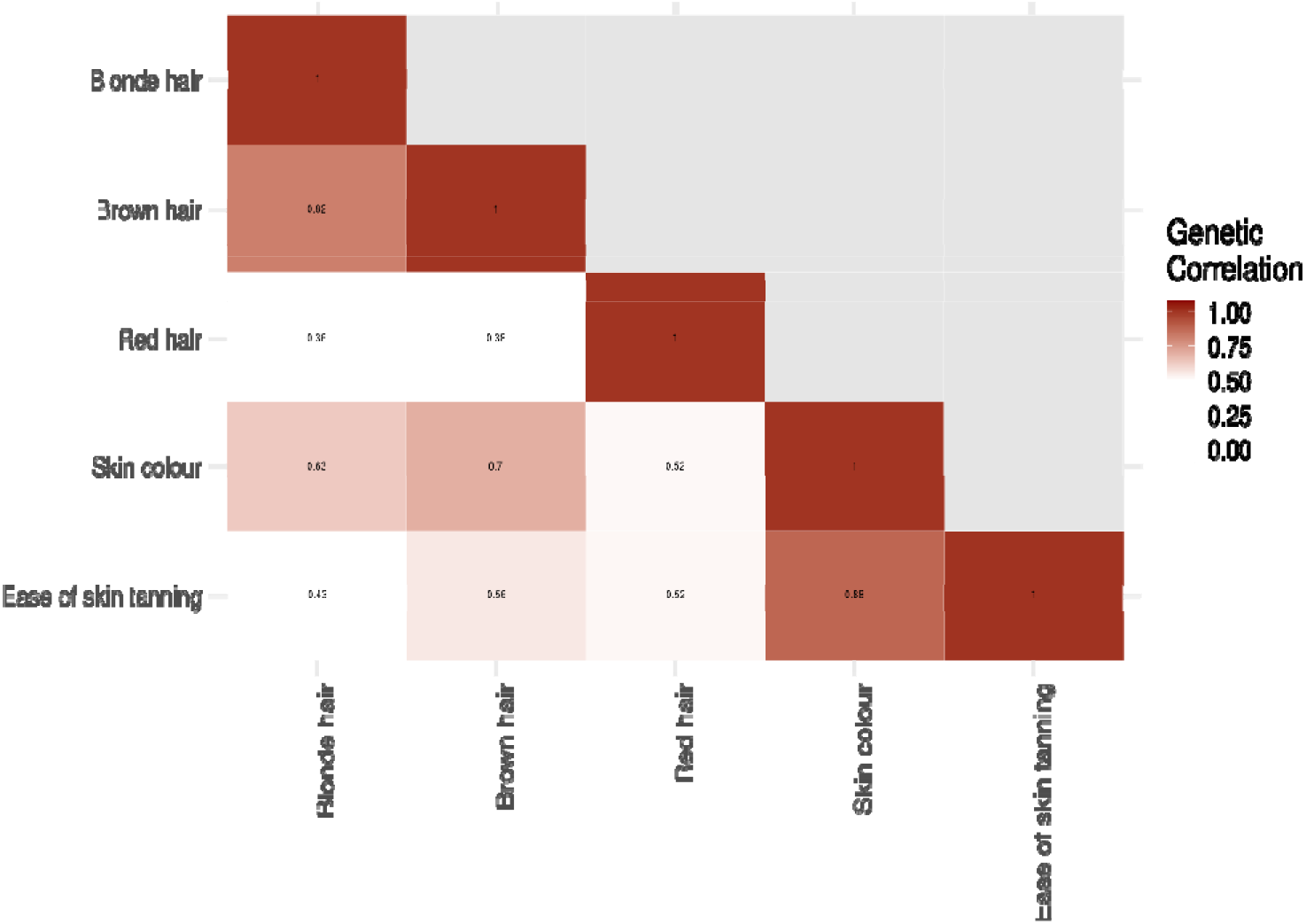
Correlation between traits. Genetic correlation between hair colour (red and blonde), skin colour and ease of skin tanning.

We identified candidate genes unique to each trait, and those in overlapping categories (Supplementary Table 19) and analysed their functional annotations using the DAVID bioinformatics tool. Unsurprisingly, the 13 candidates common to all four traits were significantly enriched for GO Terms “melanin biosynthetic process” (P= 3.7 × 10^−9^, Benjamini-Hochberg correction) and “melansomal membrane” (P=4.3 × 10^−4^, Benjamini-Hochberg correction). Furthermore, those in the three categories excluding red hair were also significantly enriched for melanosome/melanosomal membrane. Other trait categories did not show significant enrichments apart from the blonde-only category which was enriched for genes involved in hair follicle development (P=0.075) and for keratin genes (P=0.16). Neither of these gene categories are found associated with any other pigmentary trait.

## Discussion

Employing DISSECT as the tool to analyse genome wide associations in UK Biobank has clear advantages. It has allowed the inclusion of more than 100,000 individuals previously excluded due to family relatedness. As a consequence the increase in power has resulted in an increased number of SNPs associated with pigmentary traits.

We have increased by more than threefold the number of genes associated with red hair, from 6 in our previous analysis to 21. Among these are established pigmentation genes, *MITF, SLC45A2, TYR* and *OCA2* as well as the three components of the MC1R signalling pathway, *MC1R, ASIP* and *POMC*. One gene noted in several studies of red hair, *PKHD1*, has no known function in pigmentation. However, in this study we find associations with SNPs lying between *PKHD1* and the adjacent gene, *TFAP2B. TFAP2* paralogues are involved in pigmentation^28^, and it may be that it is this gene, rather than *PKHD1*, that is the causative gene in red hair colour. Our TWAS finds 7 of the associated genes have expression variation, including *MC1R*, even though coding variation can almost entirely account for the association. In contrast, a very strong candidate for a causal gene, *POMC*, does not show any associated coding variation and we assume the causal association is via expression yet it does not appear in the TWAS, highlighting limitations in the analysis.

The candidate genes associated only with blonde hair colour are enriched for hair follicle development and for keratin genes; there are 5 keratin genes in this trait, and none in any other. We have previously noted enrichment for these gene categories^8^ and speculated the association may be due to the interactions between keratinocytes and melanocytes affecting melanin transfer, or that the structure of the hair might affect its pigmentation. As skin colour does not show the same associations with keratinocyte genes, we suggest this indicates that the latter is true. Shape or thickness of the hair can affect the deposition of pigment or the perception of the pigmentation within the hair shaft. There are no melanosome/melanosomal membrane genes associated with red hair, unlike the other pigmentary characteristics. This may reflect that red/non-red variation is due to the type of melanin made (phaeomelanin vs eumelanin) and not to its packaging or delivery.

We also previously noted that female Biobank participants were more likely to self-report having red or blonde hair and paler skin than male participants. We have extended this analysis and identified five loci, *MC1R, ASIP, OCA2, IRF4* and *KITL*, at which genetic variants have a sexually dimorphic effect. What mechanism underlies the dimorphism remains to be seen, but sex hormone levels within sexes play only a minor, if any, role. A recent report^29^ finds that, within UK Biobank, melanoma incidence in men is positively correlated with testosterone levels. Our observation that men with higher testosterone levels tan less well, at least by self-report, may be relevant.

It is widely believed that the distribution of red hair in the UK is correlated with geographical origin. We show that this is in fact true, that the frequency of red hair, pale skin and poor tanning ability in UK Biobank participants is higher in the north and west of the country. In addition, we find that blonde hair is more common in those born in the east of the country. Applying PRS to these traits rather than self-reported phenotype validates these observations. Furthermore, self-reported red hair shows a stronger north-south gradient than the PRS, suggesting that individuals in the north are more likely to report red hair than those in the south.

Overall, the power of increased sample size has resulted in many more associations with pigmentary phenotypes. We nevertheless have not explained all variation; analysis of whole genome sequences of UK Biobank participants may discover more.

## MATERIALS AND METHODS

### GWAS European individuals

Study individuals were derived from the UK Biobank cohort that consists of 502,655 individuals aged between 40 and 69 years at recruitment, ascertained from 22 centres across the UK between 2006 and 2010. The study was approved by the National Research Ethics Committee, reference 11/NW/0382, and informed consent was obtained from all participants as part of the recruitment and assessment process. This particular study was approved by UK Biobank, assigned study number 7206.

We performed GWAS, using array-genotyped and imputed SNPs, on 452,264 European origin individuals from UK Biobank, which were used by the Geneatlas analysis^30^. The analysis was run with DISSECT^20^ which uses Mixed Linear Models to account for related individuals.

Quality control was applied to UK Biobank SNPs, removing SNPs with Hardy-Weinberg equilibrium <10^−50^, imputation score < 0.9 and MAF <0.01. The number of SNPs included in the analysis after QC is 9,122,998.

Hair colour phenotype was divided into two binary phenotypes: blonde hair colour (53,100 cases and 399,164 controls) and red hair colour (21,451 cases and 430,813 controls). In the red hair analysis, individuals with blonde hair or unknown hair colour were imputed to the mean value for hair colour and in the blonde hair analysis, individuals with red hair or unknown hair colour were imputed to the mean hair colour.

Skin colour is defined as a numerical phenotype with different levels 1=very fair, 2=fair, 2=light olive, 4= dark olive 5=brown and 6=black. Ease of tanning is also a numerical phenotype with 1=very tanned, 2=moderately tanned, 3=mildly or occasionally tanned, 4=never tan, 5=only burns. Table 1-4 include phenotypes for all four pigmentation traits in European origin UK Biobank individuals.

Manhattan plots for the pigmentation traits GWAS were plotted using the qqman R package. To obtain the lead SNPs for each traits, each GWAS was clumped using plink 1.9^31^ and parameters p1=5×10^−8^, p2=0.01 and r2=0.1 clump-kb=10000, Then, if the lead SNP was inside a gene, the gene was selected as the closest gene for the analysis. Otherwise the two closest genes were selected, unless it was more than 500 Kb apart from the lead SNP, in which case no gene was selected.

### Polygenic Risk Scores

Polygenic Risk Scores (PRS) for each trait were calculated using the clumped SNPs and glmnet R package ^32^. A lasso model was calculated to obtain the weights for each SNP and trait with unrelated individuals from white British ancestry. Then scores for each European individual and trait were calculated using plink 1.9. These PRS were used to calculate correlation between risk and different phenotypes.

### Whole-Exome Sequencing data

Whole-Exome Sequencing (WES) data for 200,643 UK Biobank individuals was made available in November 2020. Population bed files were used to extract the MC1R exome for all individuals and using plink 1.9. Synonymous mutations were removed from the data to keep only missense variants, insertions and deletions and then the number of individuals with 0, 1, 2 and 3 MC1R variants was calculated.

### Sex differences

UK Biobank European individuals were segregated by sex, with 206,729 men and 245,432 women. Sex-specific GWAS for hair colour (blonde and red), skin colour, and ease of tanning were performed using dissect^20^. LD score regression^24^ was used to calculate the genetic correlation between the sex-specific GWAS.

Hormone data was obtained from UK Biobank. Estradiol data is available for 68,880 European individuals (50,748 females and 18,052 males) and testosterone data for 390,585 (195,394 males and 195,191 females). Using these individuals we calculated the correlation between hormone levels and pigmentation traits, stratified by sex. We also used the calculated PRS to calculate the correlation between hormone levels and pigmentation PRS.

Information about menopause at the age of recruitment was also retrieved from UK Biobank. There are 55,736 European women with data that had not undergone menopause and had not had a hysterectomy at the age of recruitment, from them 26,426 had estradiol measurements. We repeated the correlations between estradiol levels and pigmentation traits using these individuals.

### Geographical correlation

Unrelated white British individuals (n=343,234) were used for the geographical analysis of the data. Coordinates of place of birth were extracted from UK Biobank and then calculated polygenic risk scores for the unrelated white British were used to calculate the correlation between PRS and the geographical coordinates (North-South and East-West separately).

### Transcriptome-wide association studies

RNA-seq and genotype data for 106 melanocytes were obtained from dbgap (accession number phs001500.v1.p1). RNA-seq Fastq files were aligned to ghrc37 using STAR^33^ and raw counts were calculated using featurecounts^34^. Then, TMM values for each gene were calculated using the edgeR package^35^, and a total of 16,536 genes had expression different from 0. TPM values for these 16,536 genes were also calculated using Stringtie^36^.

Genotype data from the VCF files was extracted for ±500Kb around each gene using plink^31^, and the heritability of gene expression explained by the SNPs in the region around the gene was calculated using LDscore^24^. For those genes with significant heritability (p-val<0.05), we calculated blup, elastic net, top1 and lasso models for gene expression using the Fusion R package^21^, and used a 5-fold cross validation to estimate the r^2^.

Pre-computed models of gene expression for GTEx v7 data for sun-exposed skin and not sun exposed skin were downloaded from Fusion (http://gusevlab.org/projects/fusion/#reference-functional-data).

Transcriptome-wide association studies (TWAS) for the calculated melanocyte models, sun exposed skin and not sun exposed skin using the pre-calculated models from GTExv7 were calculated using the FUSION package. Bonferroni correction was applied to gene p-values in order to find significant genes associated to each trait.

FOCUS^37^ was used to calculate the posterior probability of causality for genes in chromosome 16 for the three gene expression datasets.

## Acknowledgements

This study was funded by the UK Medical Research Council, the UK Biotechnology and Biological Sciences Research Council and the Intramural Research Program of the US National Institutes of Health via the National Human Genome Research Institute and the National Cancer Institute. The content of this publication does not necessarily reflect the views or policies of the US Department of Health and Human Services, nor does mention of trade names, commercial products, or organizations imply endorsement by the US government.

## Supplementary Information

### Supplementary Figures

**Supplementary Figure 1**

Comparison of P values of previous GWAS of unrelated white British UK Biobank participants performed using PLINK logistic regression with the present analysis of all Europeans with a linear mixed model using DISSECT. a) red hair, b) blonde hair

**Supplementary Figure 2**

Complexity of *MC1R* region. A) correlation of skin gene expression from GTex within all the genes in the MC1R region, defined as the overlapping genes with significant SNPs around MC1R and genes highlighted by TWAS analysis. B) LD within SNPs in the region around these genes.

**<u>Supplementary Figure 3</u>**

Hormone levels by sex. a) Estradiol levels from males in pink and females in blue for UK Biobank European participants and b) Testosterone levels from males in pink and females in blue for UK Biobank European participants.

**<u>Supplementary Figure 4</u>**

Logarithm of p-value for significant genes in chromosome 16 for the red hair TWAS with sun-exposed skin. Size of the points indicate posterior probability of being the causal gene calculated with FOCUS.

### Supplementary Tables

**Supplementary Table 1**

Independently associated SNPs in GWAS of pigmentary phenotypes in UK Biobank

**Supplementary Table 2**

Candidate genes for pigmentary phenotype associations in UK Biobank

**Supplementary Table 3**

Distribution of hair colour for 200K individuals with WES data available

**Supplementary Table 4**

Number of MC1R variants per individual with red hair

**Supplementary Table 5**

Red hair associated variants of MC1R, from exome sequencing. MAF: minor allele frequency in this cohort, MAC: minor allele count.

**Supplementary Table 6**

Genetic correlation between sexes for the different pigmentation traits

**Supplementary Table 7**

SNPs with significant differences in a sex-specific red hair GWAS in UK Biobank European population. Data shows chromosome, position, summary statistics for the male and female gwas and p-value of the difference.

**Supplementary Table 8**

SNPs with significant differences in a sex-specific blonde hair GWAS in UK Biobank European population. Data shows chromosome, position, summary statistics for the male and female gwas and p-value of the difference.

**Supplementary Table 9**

SNPs with significant differences in a sex-specific skin colour GWAS in UK Biobank European population. Data shows chromosome, position, summary statistics for the male and female gwas and p-value of the difference.

**Supplementary Table 10**

SNPs with significant differences in a sex-specific ease of skin tanning GWAS in UK Biobank European population. Data shows chromosome, position, summary statistics for the male and female gwas and p-value of the difference.

**Supplementary Table 11**

Correlation between hormone levels and pigmentation stratified by sex for UK Biobank European individuals

**Supplementary Table 12**

Correlation between hormone levels and a polygenic risk score for pigmentation traits stratified by sex in UK Biobank European individuals

**Supplementary Table 13**

Correlation between place of birth and pigmentation phenotypes using a polygenic risk score and the phenotypic self-reported data from UK Biobank

**Supplementary Table 14**

Red hair TWAS summary statistics for sun exposed skin, not sun exposed skin from GTEx and melanocytes transcriptome data

**Supplementary Table 15**

Blonde hair TWAS summary statistics for sun exposed skin, not sun exposed skin from GTEx and melanocyte transcriptome data

**Supplementary Table 16**

Skin colour TWAs summary statistics for sun exposed skin, not sun exposed skin from GTEx and melanocyte transcriptome data

**Supplementary Table 17**

Ease of tanning TWAS summary statistics for sun exposed skin, not sun exposed skin from GTEx and melanocyte transcriptome data

**Supplementary Table 18**

Overlaps between GWAS and TWAS candidate genes

**Supplementary Table 19**

Candidate genes in common between pigmentary traits

