## Supplementary figures and images for "Expanded Analysis of Pigmentation Genetics in UK Biobank"

### Supplementary Figure 1

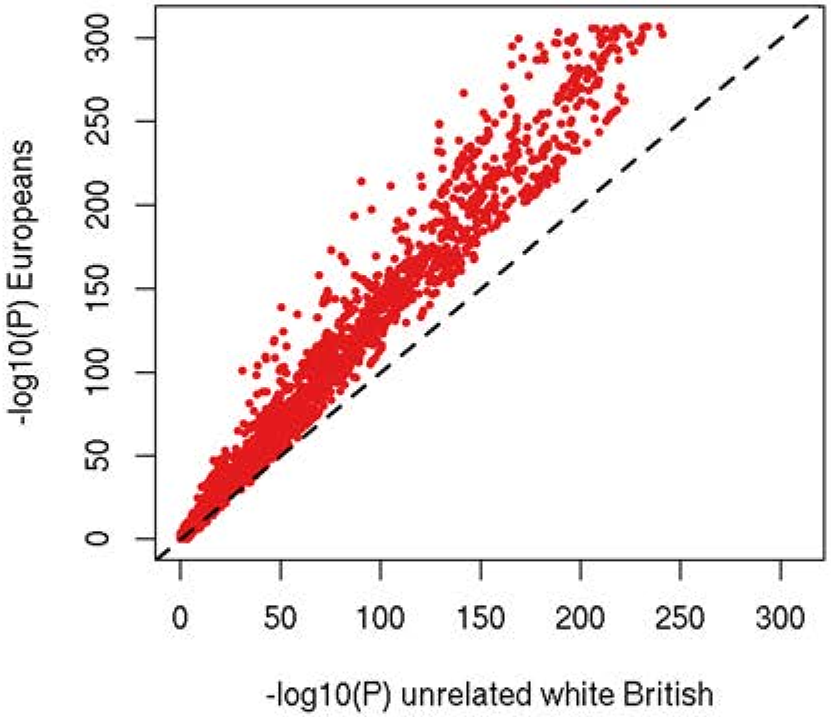

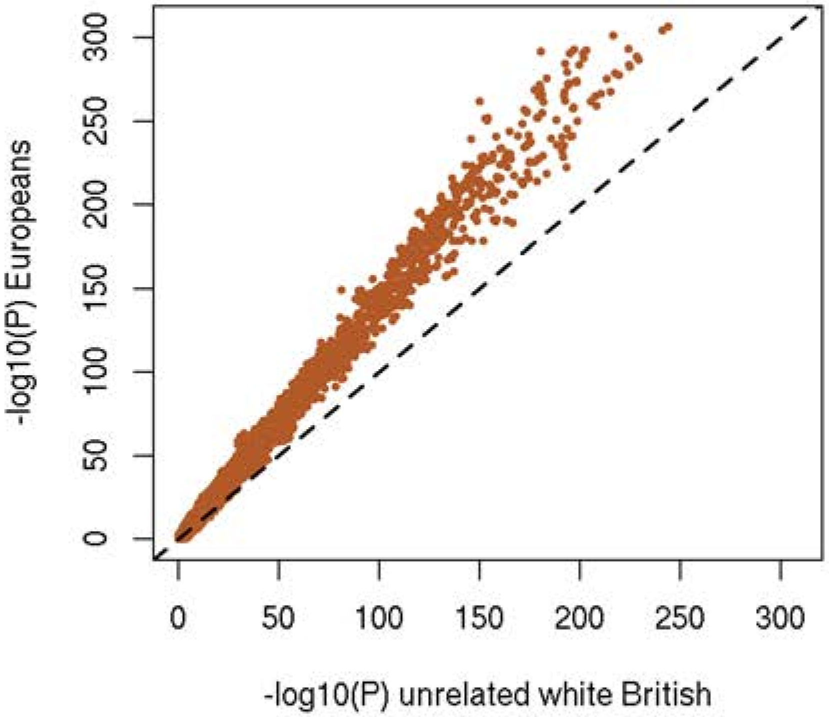

### Supplementary Figure 2

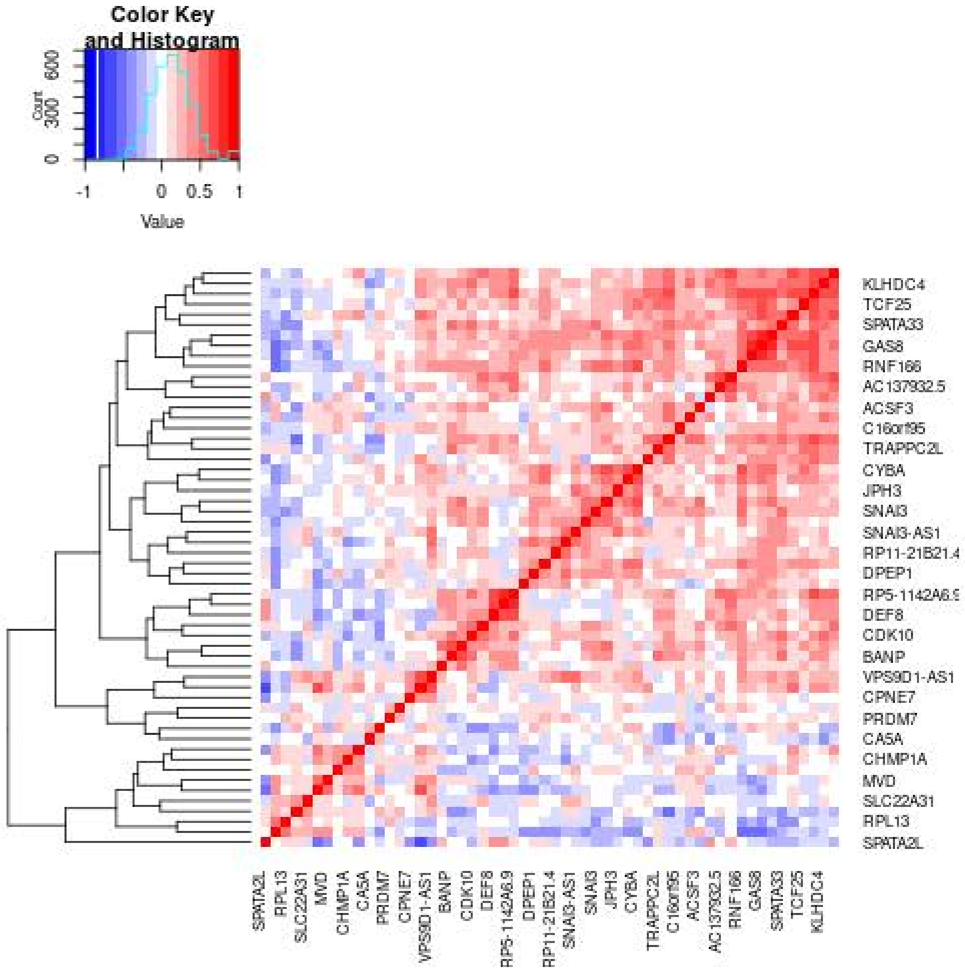

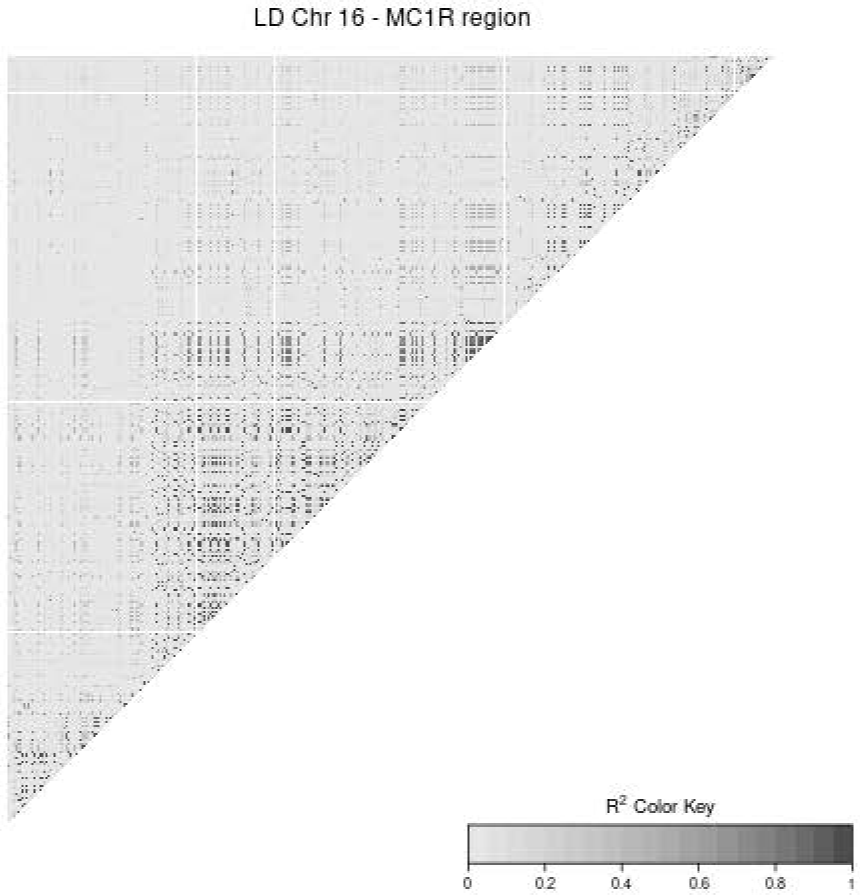

### Supplementary Figure 3

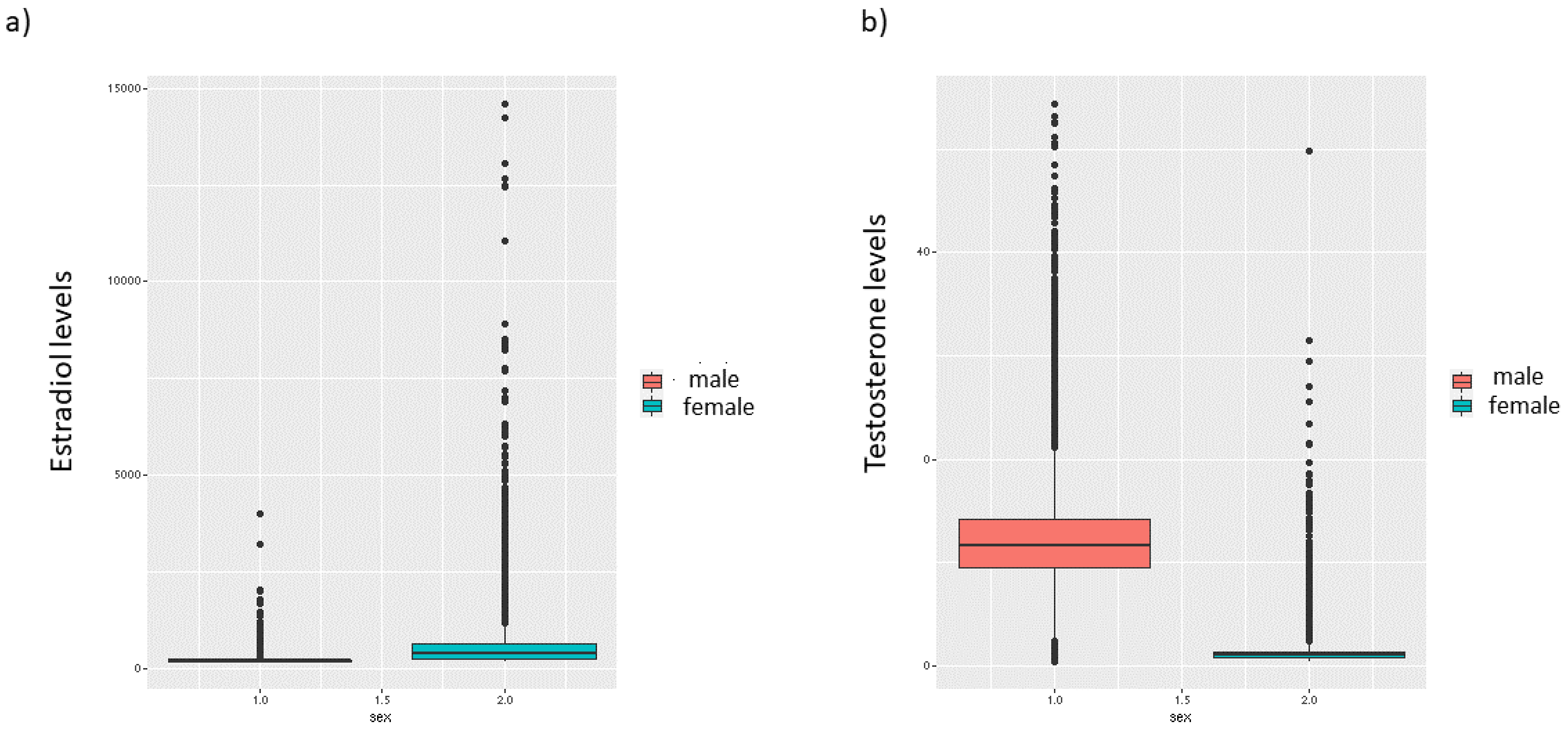

### Supplementary Figure 4

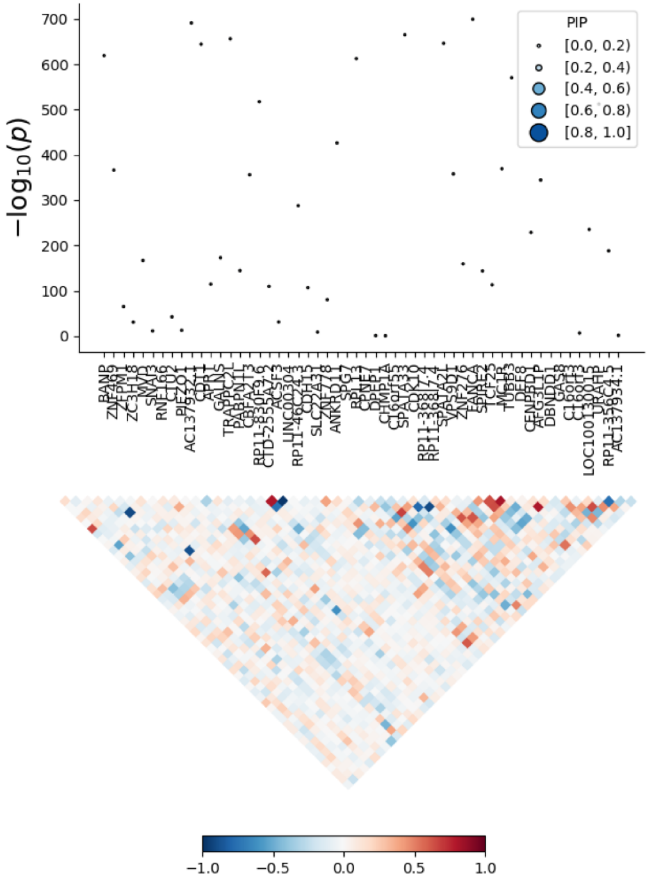
