## Supplementary Tables 3, 4, 6, 13 for "Expanded Analysis of Pigmentation Genetics in UK Biobank"

Supplementary Table 3: Distribution of hair colour for 200K individuals with WES data available

| **Hair colour** | **Number of individuals** | **Percentage** |
| --- | --- | --- |
| Blonde | 21182 | 10.55% |
| Red | 8496 | 4.23% |
| Light Brown | 76582 | 38.16% |
| Dark Brown | 74924 | 37.34% |
| Black | 16522 | 8.23% |
| NA | 2937 | 1.46% |

Supplementary table 4: Number of MC1R variants per individual with red hair

| MC1R coding variants | Individuals | Percentage of red hair |
| --- | --- | --- |
| 0 | 22 | 0.2% |
| 1 | 207 | 2.4% |
| 2 | 8496 | 96.2% |
| 3 | 97 | 1.14% |

Supplementary table 6: Genetic correlation between sexes for the different pigmentation traits

| **Trait** | **Genetic correlation male/female** |
| --- | --- |
| Blonde | 1.02(0.03) |
| Red | 0.950(0.06) |
| Skin colour | 0.967(0.02) |
| Ease of tanning | 0.956(0.02) |

Supplementary table 13: Correlation between place of birth and pigmentation phenotypes using a polygenic risk score and the phenotypic self-reported data from UK Biobank

| **trait** | **N-S PRS** | **N-S pheno** | **E-W PRS** | **E-W pheno** |
| --- | --- | --- | --- | --- |
| Red | 0.014 | 0.027 | -0.012 | -0.016 |
| Blonde | -0.015 | 0.0029 (NS) | 0.024 | 0.010 |
| Skin colour | -0.030 | -0.028 | 0.02 | 0.0044 |
| Ease of tanning | 0.028 | 0.031 | -0.02 | -0.014 |
