## Supplementary Table 11 for "Expanded Analysis of Pigmentation Genetics in UK Biobank"

Supplementary Table 11. Correlation between hormone levels and Pigmentation traits (self-reported) by sex. Significance after multiple test correction is indicated with *

| **Hormone** | **Blonde (Pheno)** | **Red**  **(Pheno)** | **Male Blonde (Pheno)** | **Female Blonde**  **(Pheno)** | **Male Red (Pheno)** | **Female Red (Pheno)** | **Skin colour (Pheno)** | **Ease of tanning (Pheno)** | **Male Skin colour** | **Female Skin colour**  **(Pheno)** | **Male Ease of tanning**  **(Pheno)** | **Female Ease of tanning** |
| --- | --- | --- | --- | --- | --- | --- | --- | --- | --- | --- | --- | --- |
| Testosterone | -0.042* | -0.036* | -0.002 | 0.0007 | 0.0002 | 0.002 | 0.019* | -0.12* | 0.004 | 0.003 | -0.017* | 0.002 |
| Estradiol | 0.008 | 0.007 | 0.0008 | -0.006 | -0.009 | -0.004 | -0.001 | 0.006 | 0.007 | 0.003 | 0.012 | -0.0008 |
