## Supplementary Table 12 for "Expanded Analysis of Pigmentation Genetics in UK Biobank"

Supplementary Table 12 Correlation between hormone levels and Polygenic risk scores for pigmentation traits

| **Hormone** | **Blonde** | **Red** | **Male Blonde** | **Female Blonde** | **Male Red** | **Female Red** | **Skin colour** | **Ease of tanning** | **Male Skin colour** | **Female Skin colour** | **Male Ease of tanning** | **Female Ease of tanning** |
| --- | --- | --- | --- | --- | --- | --- | --- | --- | --- | --- | --- | --- |
| Testosterone | -0.001 | -0.0002 | -0.007 | -0.002 | -0.003 | 0.005 | -0.002 | -0.0005 | 4e-5 | 0.002 | -0.003 | 0.004 |
| Estradiol | 0.0002 | 0.0004 | 0.002 | -0.002 | -0.001 | -0.002 | 0.002 | -0.003 | -0.002 | 0.002 | 0.003 | -0.003 |
